# Brain Injury Knowledge Ontology (BIKO) for traumatic brain injury: Formalizing concepts and methods used in translational traumatic brain injury research

**DOI:** 10.1101/2023.10.29.564650

**Authors:** Monique C. Surles-Zeigler, Troy Sincomb, C Edward Dixon, Fahim Imam, Tom Gillespie, Jeffrey S. Grethe, Adam R. Ferguson, Maryann E. Martone

**Affiliations:** University of California-San Diego, Center for Biological Research, San Diego, California; University of California-San Francisco, Neurological Surgery, San Francisco, California Ed Dixon

## Abstract

Traumatic brain injury (TBI) is an insult to the brain resulting from an external force and is a significant cause of morbidity and mortality in the United States. No effective clinical therapeutics currently exist for this injury. Although several therapies and procedures have been deemed successful for TBI treatment in preclinical research studies, they have yet to be translated into human patients. These discouraging results have left many scientists questioning the role of animal models in drug discovery for TBI.

One major hurdle in translating the knowledge obtained in the laboratory to the clinic is the methodological variance across these studies. This variance can hinder the ability to draw conclusions from conflicting studies and aggregate data across various research studies, which ultimately impedes the ability to aggregate data across these studies. Therefore, addressing this variance is crucial for bridging the gap between the laboratory and the clinic. The increasing volume of papers and associated data being published every day makes this hurdle even more difficult to overcome. The initial steps to address these knowledge gaps are identifying these studies and creating a shared knowledge framework for mapping their terminology. We are developing the Brain Injury Knowledge Ontology (BIKO) to create a standardized model to describe methods and outcome measures used within preclinical and clinical TBI therapy studies to facilitate comparison across studies and models. The first version of BIKO focuses on modeling the major preclinical TBI models, e.g., Controlled Cortical Impact Model, Fluid Percussion Model, and Weight-Drop Model), major neurological injuries related to these models and their relationship to clinical pathophysiology. We show how BIKO provides a machine-readable way to represent the methodologies used in TBI therapeutic studies to compare models across clinically relevant features.

## Introduction

Traumatic brain injury (TBI) is a significant health issue predicted to be a leading cause of disease burden globally by 2030 (Mathers and Loncar 2006). In the United States alone, TBI affects an estimated 1.5 million people each year, claiming the lives of approximately 50,000 people annually (Korley et al. 2016) and leaving the remainder often with significant disabilities. A TBI is caused by an insult to the brain by an external force originating from an injury event (e.g., car accident, fall, motor vehicle accident, blast). TBI is a very complex injury involving two distinct stages: primary and secondary. Primary injury is the initial injury to brain tissue caused by mechanical disruption at the time of injury, activating many biological pathways.

Secondary injury represents the ongoing injury initiated by the primary injury that can last for minutes, hours, or years. Due to the multitude of activated pathways, TBI is also a risk factor for other neurological diseases that decrease life quality in these patients, including epilepsy (Salazar et al. 1985; Jennett and Lewin 1960; Coulter et al. 1996), sleep disorders (Wilde et al. 2007; Castriotta et al. 2007; Verma, Anand, and Verma 2007), stroke (Liu et al. 2017; Burke et al. 2013), Alzheimer dementia (Green et al. 2020; Masel and DeWitt 2010), and Parkinson disease (Delic et al. 2020; Masel and DeWitt 2010).

Due to the complexity and potentially long time scale of the second stage of TBI, it has been challenging to create successful treatments. In 2019 alone, the National Institutes of Health spent an estimated 133 million dollars on TBI research, with approximately $12 million on preclinical therapeutic studies (National Institutes of Health 2020). The majority of our current understanding of how the brain responds to TBI and potential therapies comes from animal studies. While many preclinical studies have successfully reduced neuronal cell death following injury, these results have not successfully translated to human patients. In hopes of understanding this gap in translation, several groups have examined potential reasons why these therapies have not been successfully translated to clinical studies. Their conclusions identified insufficient preclinical sample size, functional outcome measurements, and biomarkers used, or potential interactions between therapy and anesthetic used in preclinical animals (Kabadi and Faden 2014; Loane and Faden 2010; Loane, Stoica, and Faden 2015). As the pre-clinical literature is vast, comprising several thousand published studies, it is difficult for a human to compare these studies to gain an understanding of how methodological variance impacts outcomes.

An initial step towards addressing the translation of knowledge within and to the clinic is to provide a structured framework for standardizing terminology and relationships between TBI concepts within the TBI field - an ontology. There is currently no ontology to organize and aid TBI translational research. In simple terms, an ontology is a map or framework that assists computers in processing and categorizing information about a specific domain to facilitate scientific discovery and data integration. Ontologies have been successful in organizing a field, with efforts like Gene Ontology to align gene products over multiple species and biological, molecular, and cellular function (Botstein et al. 2000; Gene Ontology Consortium 2001; “[PDF] A Short Study on the Success of the Gene Ontology ..,” n.d.). The use of GO as an annotation standard has resulted in large knowledge bases that allow scientists around the world to identify genes and functions across species and create gene networks predicting the relationships between genes (The Gene Ontology Consortium 2019; Pirooznia et al. 2012; Montojo et al. 2014; Maere, Heymans, and Kuiper 2005).

The Brain Injury Knowledge Ontology (BIKO) is designed to provide a standardized method to assert experimental design parameters (injury models and methods) and outcome measures used in preclinical and clinical TBI therapy studies. We describe here the first release of BIKO, which is focused on modeling the relationship between clinical and preclinical TBI, the major preclinical TBI models and assessments used to assess pathological, cognitive, and behavioral outcomes of TBI.

## Methods

BIKO-TBI (RRID:SCR_024628) was developed using the OWL 2 Web Ontology Language and RDF Schema. We used WebProtege (RRID: SCR_024627), Protege (RRID: SCR_003299, version 5.5.0), and the Python RDFlib (RRID: SCR_024629, version 7.0.0) ontology development tools to structure and build the ontology and Hermit (RRID:SCR_016006), version 1.4.3.456) was used for the reasoning. Each BIKO entity is assigned both a compact identifier of the form BIKO:###### and has an Internationalized Resource Identifier (IRI) - https://www.bikotbi.org/uri/{compact_identifier}. Relevant terms were imported from pre-existing ontologies identified and accessed through Bioportal (Musen et al. 2008; Salvadores et al. 2013; Whetzel et al. 2011) or Ontology Lookup Service (Côté et al. 2006, 2008; Harrow et al. 2018; Jupp et al. 2015).

BIKO was designed in accordance with best practices espoused by the Open Biological and Biomedical Ontology (OBO)(The OBO Foundry 2021) and FAIR principles (Wilkinson et al. 2016; Guizzardi 2020; Garijo and Poveda-Villalón 2020). Such practices include providing open access to BIKO (Final Github version #), using a common formal language (OWL) to enhance interoperability, and adopting a modular design. We used components from the Basic Formal Ontology (BFO) upper ontology to promote interoperability with similar ontologies, principally the Regenebase ontology for pre-clinical spinal cord injury (Callahan et al. 2016)(Figure 1), Alzheimer’s Disease Ontology (ADO,(Malhotra et al. 2014)) and Epilepsy Ontology (EPIO, (Sargsyan et al. 2023)). Thus, the top structure of BIKO-TBI comprises 4 classes: attribute, information content entity, material entity, and process, a subset of the upper-level RegenBase ontology (Fig 1). BIKO is accessible in multiple formats (.owl, .ttl, .rdf), including as a single file. The merged version of BIKO underwent manual curation to integrate ontology slims into the core structure of BIKO.

**Figure 1.**
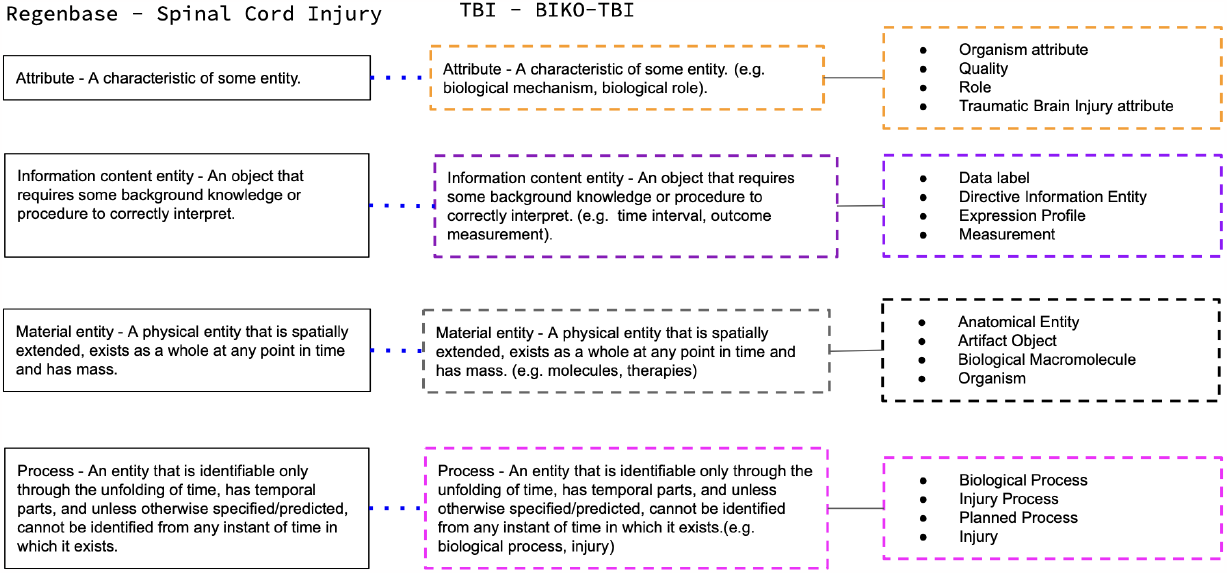
Top Level Ontology Schema. BIKO-TBI was developed to align with similar ontologies like Regenbase, Alzheimer’s Disease Ontology (ADO), and Epilepsy Ontology (EPIO). This figure illustrates the alignment of the top-level schema across the two ontologies and the associated BIKO subclasses, with BIKO containing the same upper-level classes, allowing for interoperability across the two neurotrauma knowledge bases.

The structure and content of BIKO are constructed from multiple knowledge sources to increase reuse and interoperability, including:

1. pre-existing terms within community ontologies: Relevant terms and classes from community ontologies were identified through BioPortal. All mapped terms were mapped by one of two methods. For branches containing less than 10 terms, they were linked to their IRIs using the rdfs:SeeAlso (“RDF 1.2 Schema” n.d.) annotation property. For branches containing 10 or more terms, a second method involved creating a slim version, a condensed subset of an external ontology branch. These slim modules underwent manual curation and were integrated into the core structure of the merged version of BIKO. Branches of Foundational Model of Anatomy (FMA, Brain Segment - FMA:55676, version 5.0.0) and National Cancer Institute Thesaurus (NCIT, Organism - NCIT:C14250, version 23.09d). Future versions will include a merged slim from Uber-anatomy ontology (UBERON). Using these two mapping approaches, we aimed to streamline BIKO’s management, maintenance, and organization.
2. the scientific literature: Major classes and entities related to clinical and preclinical TBI were derived from reviews and primary experimental papers on TBI identified through multiple manually curated PubMed searches related to preclinical models of TBI.
3. TBI domain experts: Domain experts from the PRECISE-TBI initiative provided expert knowledge and feedback on BIKO structure and content. PRECISE-TBI (PREClinical Interagency reSearch resourcE for Traumatic Brain Injury) is an interagency (VA, NIH, DoD) project designed to elevate “rigor, reproducibility, transparency” in preclinical TBI research to improve translation to the clinic.
4. PRECISE Model Catalog (RRID:SCR_024626) and pre-clinical TBI CDE’s:

BIKO-TBI is an application ontology and an information system that will be used by downstream tools such as knowledge bases and other computational methods to integrate and analyze TBI data. As such, we aligned preclinical TBI CDEs (Smith et al. 2015; Hicks et al. 2013) to BIKO classes and added sample metadata from 42 studies listed in the PRECISE-TBI model catalog. These studies were considered instances of the major pre-clinical model types. The Model Catalog is a queryable online knowledge base to discover and compare preclinical TBI models and link them to additional information such as protocols and datasets model literature metadata (scicrunch.org/precise-tbi, precise-tbi.org).

### Evaluation of BIKO-TBI

To ensure the methodological rigor of the model and the associated system performance, a set of competency questions was developed to guide development and check accuracy. The queries listed below were developed to the current version’s performance.

- CQ1: Find all preclinical models of open head injury.
- CQ2: Find all open head injury models with predominant focal damage.
- CQ3: Show all preclinical assessments of motor function.

## Results

Overview of BIKO V1.0: Figure 2 provides an overview of the contents of BIKO-TBI V1.0. BIKO-TBI (Fig 2) contains 15543 terms with imports from FMA and NCIT. Most of the terms in BIKO-TBI v.1.0 contains associated synonyms or abbreviations, definition, definition source, and external identifier.

**Figure 2.**
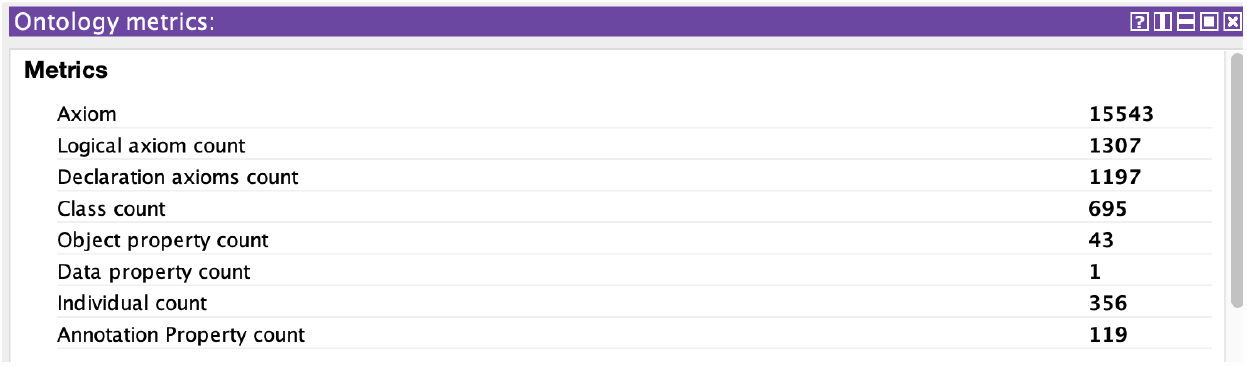
Current status of BIKO as of October 2023. A pre-release of the ontology .owl file is available at https://github.com/MCSZ/bikotbi.io/releases

BIKO v1 focuses on modeling both TBI and pre-clinical models of TBI. Following the modeling in RegenBase, a TBI is considered a type of injury resulting from an injury event involving a force applied to the head. The injury itself is considered a process, also following RegenBase. The advantage of modeling injuries like TBI as a process, as opposed to a continuant (something that exists at discrete time points), is that it provides the flexibility to capture the dynamic nature of the injury and allows for additional modeling of its temporal aspects.

In BIKO, injuries are classified into two types, controlled or uncontrolled, to distinguish between the injuries experienced by subjects in the laboratory versus clinical injuries sustained in the real world through events such as accidents, explosions, etc. (Fig 3).

**Figure 3.**
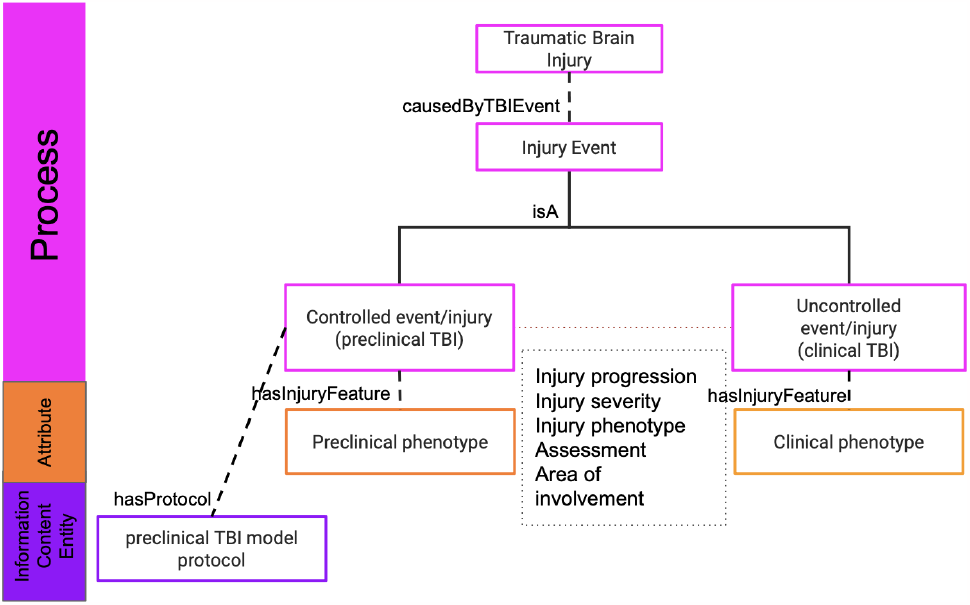
Modeling traumatic brain injury (TBI) as a process. The model distinguishes between preclinical TBI models and clinical TBI via the paradigms used to produce TBI.

### TBI Model

As BIKO was designed to facilitate the comparison of preclinical models of TBI to the clinical condition, we focused in the initial version on creating definitions of the major model types (Fig 3). Specifically, we concentrated our efforts on the 4 major TBI models: the controlled cortical impact, fluid percussion, weight drop, and blast models to make the development more manageable. Within BIKO, define a preclinical model of TBI as a process whereby a laboratory animal in a controlled laboratory environment receives a TBI according to a specification provided in a protocol. Protocols are, therefore, represented as information content entities comprising instructions, procedures, and methods created by and used in the TBI field to cause a brain injury. Each protocol records the intended mechanism of injury (e.g., fluid wave impact) and the materials and methods required, e.g., devices used to effect the injury. Protocols are then executed in a laboratory, initiating the injury event (blast, controlled cortical injury) within an animal, resulting in a TBI (Fig 4). This injury protocol is one of the distinguishing differences between a TBI occurring in the lab (controlled setting) versus a clinical TBI (uncontrolled setting).

**Figure 4.**
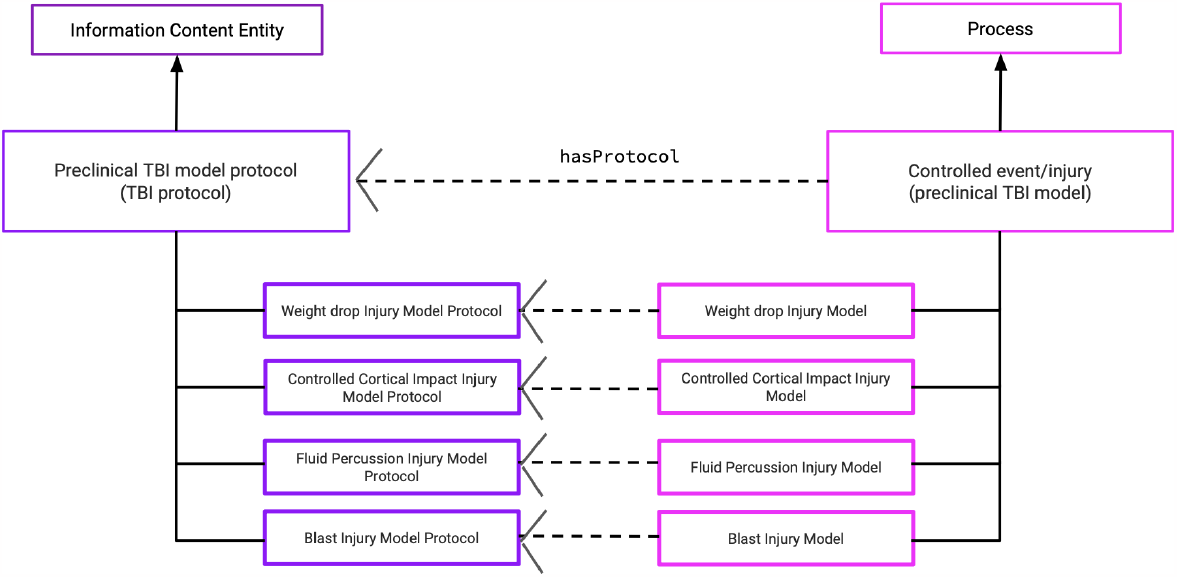
Modeling preclinical TBI protocol and the injury caused by implementing the protocol in BIKO. The figure illustrates the relationship between a controlled injury and a TBI model, where a controlled injury hasProtocol Preclinical TBI model protocol.

Execution of the standard protocol for each model results in TBIs in animals that share phenotypic features with the clinical condition, including injury type (focal vs diffuse injury), integrity of the skull (closed versus open injury), and the general area of injury (e.g., midline, lateral to bregma). To relate pre-clinical TBI to outcomes, we have also included the major types of assessments used to measure anatomical, physiological, and behavioral outcomes of TBI as types of planned processes (BIKO:000073). For the latter, we include the most common preclinical behavioral tests and relate them to the domain they are asserted to measure e.g., cognitive or motor function.

Lastly, with the significant effort being put into the development and use of CDEs for pre-clinical TBI (La Placa et al., 2021) via the PRECISE project (https://www.precise-tbi.org/) to harmonize across datasets, we have started to align the concepts in BIKO with these CDEs.

CDEs clearly distinguish between the different types of models and the associated data obtained from these preclinical models. When these CDEs are harmonized with concepts in BIKO, it enhances data interoperability, enabling data sharing across diverse injuries, and plays an important role in the development of downstream applications like knowledge bases.

### Modeling TBI injury

As noted previously, the classification of a preclinical model of traumatic brain injury as a process offers the advantage of capturing the dynamic aspects of a TBI, which evolves over time. This alignment with the real-world complexity and clinical implications of TBI is crucial. It’s important to note that the classification of a preclinical TBI model differs from that of a clinical TBI. While the preclinical model encompasses certain injury components found in the real world, it is executed in a controlled laboratory environment. TBIs are often categorized in several ways, e.g., by the mechanism of initial injury to the skull - open vs closed injury. For clinical situations, an open head injury occurs when there is some penetrating object that breaches the skull or severe skull fracture upon initial injury. In contrast, in a closed head injury, the skull is intact, e.g., a concussion. Animal models of open head injury typically consist of a portion of the skull being mechanically removed (controlled breach), e.g., by a craniotomy or craniectomy. The skull is not removed in a closed-head injury model.

In BIKO, skull integrity is modeled as a traumatic brain injury attribute in addition to impact depth, impact duration, and severity of injury. An animal who has received a surgical procedure to remove part of the skull, e.g., a craniotomy, is modeled as having a hasSkullFeature skull breach. This manual skull breach of the skull occurs after ‘some’ hasInjuryMechanism craniotomy in a preclinical TBI model.

Table 1 shows the major characteristics of the four types of preclinical models included in BIKO v1: Controlled Cortical Impact, Weight-Drop (and major variants), Blast and Fluid percussion injury models (and major variants). In the following section, we show how these models can be retrieved by query according to knowledge encoded in BIKO by working through the competency questions described in the methods section.

**Table 1.** List of the injury features associated with the preclinical model in BIKO.

| Preclinical Model | Skull Integrity | Focal/Diffuse structural damage | Site of primary injury | References |
| --- | --- | --- | --- | --- |
| Weight Drop Model |  |  |  |  |
| Feeney's Weight Drop | Manual Skull breach | Focal | Cortex | (Marmarou et al. 1994; Foda and Marmarou 1994; Dail et al. 1981; Feeney et al. 1981) |
| Shohami's Weight Drop | Non-breach skull | Predominately Focal | Global | (Chen et al. 1996; Shapira et al. 1988) |
| Marmarou's Weight Drop | Non-breach skull | Predominantly Diffuse | Global | (Marmarou et al. 1994; Foda and Marmarou 1994; Flierl et al. 2009) |
| Maryland's Weight Drop | Non-breach skull | Predominantly Diffuse | Frontal Cortex | (Kilbourne et al. 2009; Hayes et al. 1987; McIntosh et al. 1987) |
| Controlled Cortical Impact Model | Manual Skull breach | Predominantly Focal | Cortex | (Osier and Dixon 2016; Dixon et al. 1991; Lighthall 1988; Lighthall, Goshgarian, and Pinderski 1990; Hannay et al. 1999; Manley et al. 2006) |
| Blast Model | Non-breach skull | Predominantly Diffuse | Global | (Cernak et al. 1996; Goldstein et al. 2012; Bauman et al. 2009) |
| Fluid Percussion |  |  |  |  |
| Middle | Skull breach | Predominantly Diffuse | Cortex | (Hayes et al. 1987; McIntosh et al. 1987; Millen, Glauser, and Fairman 1985; Pfenninger et al. 1989; Dixon et al. 1987) |
| Lateral | Skull breach | Predominantly Focal | Cortex | (McIntosh et al. 1989; Carbonell et al. 1998) |

### Competency queries

To guide the development of BIKO-TBI v1 and to test for accuracy and completeness, we focused on 3 basic competency questions. These queries are listed below with a detailed explanation of the modeling, the expected answer, and the answer provided by the ontology.

#### Competency query 1 - Find all preclinical models of open head injury

This query was translated as “Find preclinical TBI model and isAboutTBI some ‘Open head injury’ and returned correctly only the 5 pre-clinical models which involved a breach of skull. Note that all models were asserted to be about a TBI that has skullFeature “removal of some portion of skull”. BIKO successfully traversed the hierarchy to correctly classify these as types of open head injury as shown in Fig. 5.

**Figure 5:**
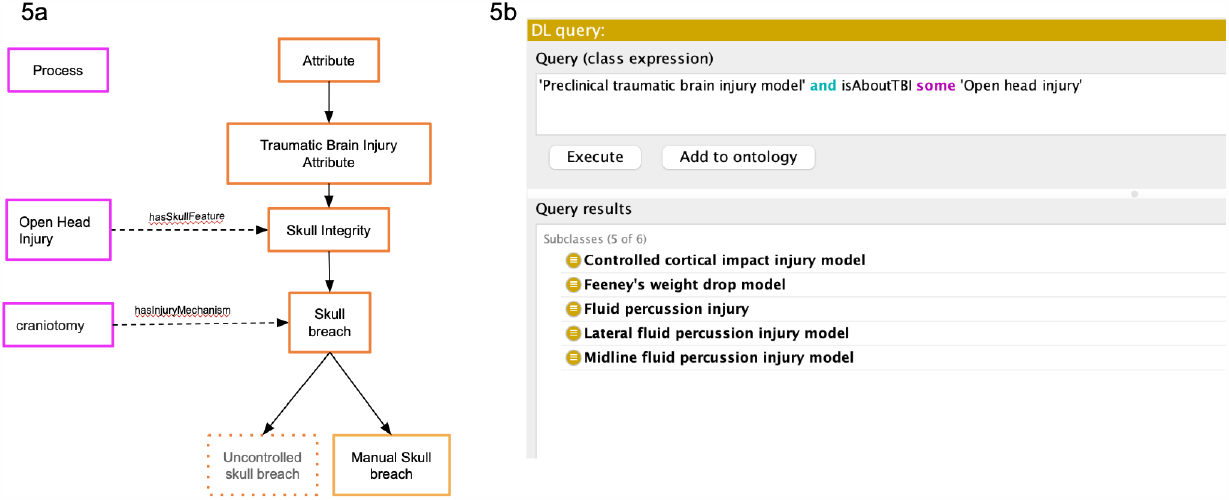
Modeling and results for Competency Question 1 (CQ1). (5b) CQ1 queries BIKO for all existing preclinical models with an open head injury. (5a) Within BIKO, a preclinical TBI is categorized as an open-head injury if there is some manual removal of the skull (craniotomy).

#### Competency query 2 - A) Find all open head injury models with predominant focal damage; B) Find all preclinical TBI models of focal damage

Clinical injuries are not solely categorized as closed and open-head injuries; but also be classified based on the extent of damage that occurred as part of the primary injury. This classification allows an injury to be categorized as focal, where the primary damage is predominantly confined to a specific region of the brain, or diffuse, where the initial damage is more widespread throughout the brain. We manually asserted TBI models as predominantly exhibiting focal or diffuse characteristics. This was carried out with the “hasPredominatelyInjury’’ relationship, including the multitude of associated injury components.

For query 2, the objective was to identify TBI models that are both open-head injury models, as per CQ1, with predominant focal damage. As expected, the query returned the expected results (Fig 6a): Feeney Weight Drop, Middle Fluid Percussion Injury, Lateral Fluid Percussion Injury, and Controlled cortical impact (Xiong, Mahmood, and Chopp 2013). Fig 6b shows the result of a related query: “Find all preclinical models of focal injury.” In this case, the Shohami weight drop model creates a closed head injury model but results in focal damage. Thus, BIKO was able to distinguish among models along the various dimensions of TBI, including the extent of the damage. These queries required no additional reasoning beyond CQ1, as focal vs diffuse is an asserted property. However, they were used to test the accuracy of the knowledge in BIKO with domain experts to ensure that models were accurately represented in this dimension.

**Figure 6:**
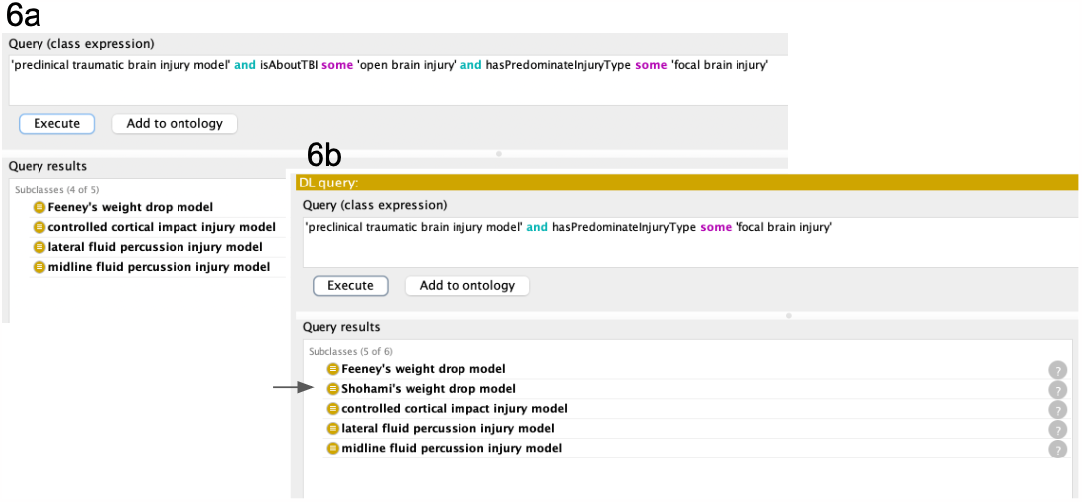
Screenshots of Protege DL query showing competency queries (CQ)2. (a) Find all closed head injury models with predominant focal damage. (b) Find all preclinical models of focal brain injury.

#### Competency query 3 - Show all preclinical assessments of motor function

As with any preclinical study of disease/injury, it is important to relate a preclinical behavioral test to the behavior it purports to measure. This connection is especially essential to relate preclinical tests to their counterparts in human studies. In this version of BIKO, we use a coarse characterization of behavioral tests as measuring predominately cognition, learning, memory, motor, sensory, or affective functions. These functions are then assigned to specific assessments via the “isAboutFunction” property (Fig 7). In future versions of BIKO, we will expand this feature to include more specific behaviors, e.g., spatial memory.

**Figure 7:**
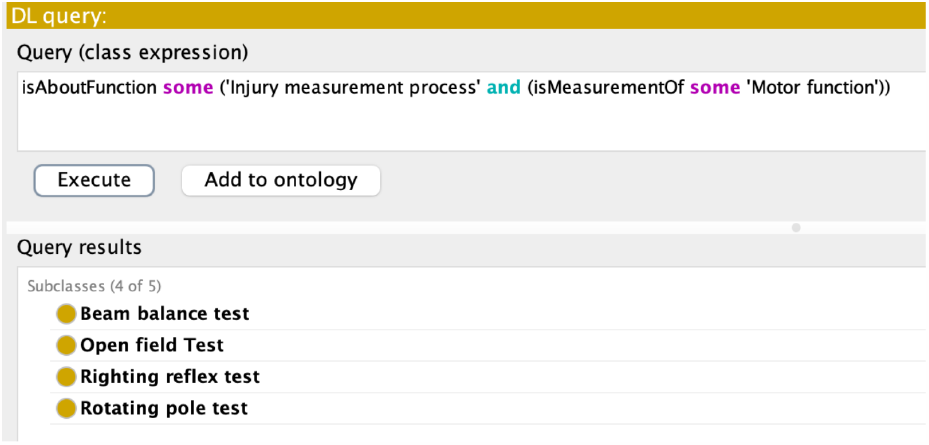
Competency queries (CQ) 3. Show all preclinical assessments that have a role in some motor function assessment.

### Use case: Applying BIKO to describe pre-clinical studies of TBI

As previously mentioned, the primary objective of BIKO is to serve as an application ontology, which implies its utility in practical scenarios. To illustrate this concept, we conducted a proof-of-concept exercise using data from a key BIKO stakeholder, PRECISE-TBI. PRECISE-TBI is a collaborative community of TBI experts dedicated to accelerating the development of therapies for TBI by increasing rigor, reproducibility, and transparency in preclinical research. The PRECISE catalog, a searchable online repository of metadata from preclinical TBI model papers in PubMed contains the four fundamental models in the current version of BIKO: Controlled Cortical Impact, Weight-Drop, Blast, and Fluid percussion injury models(Fig 8). The catalog provides a detailed search of curated preclinical TBI article metadata updated quarterly. For this exercise, we aligned the metadata from 42 research papers in the model catalog with BIKO, focusing on the citation, models, sex, species, strain, and TBI model. Subsequently, we performed Query 1 to “Find all instances of preclinical models of open head injury “Our goal was to identify studies of models referenced within the papers according to semantics encoded in BIKO. This allows us to query for clinically useful concepts such as “open head injury” that are not explicitly encoded in the Model Catalog. Using both together, we can compare attributes of these studies, e.g., sex and age of animals and devices used to study open head injury detailed in Query 1 (Fig 9).

**Figure 8:**
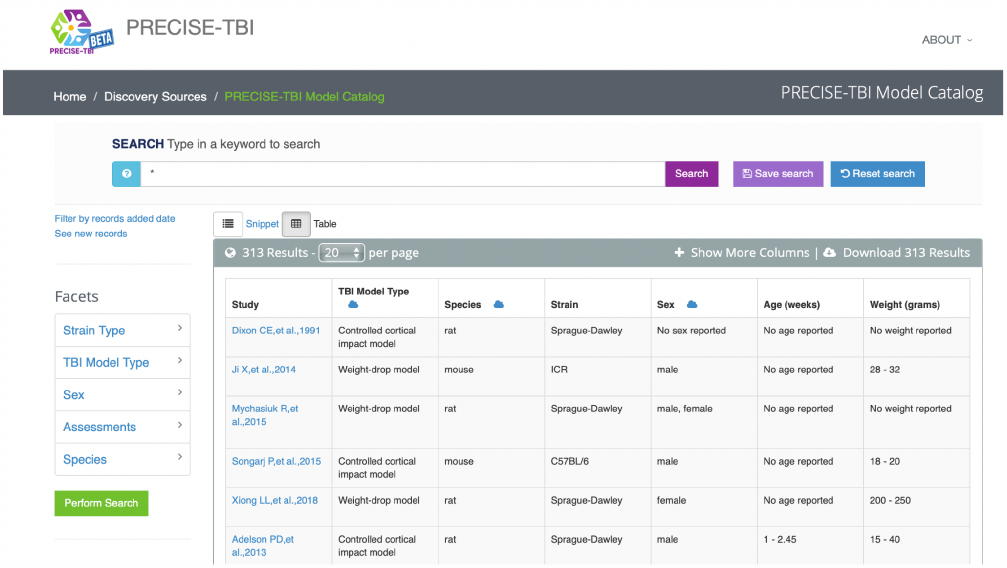
Screenshot of the PRECISE-TBI model catalog accessible at https://scicrunch.org/precise-tbi/about/model-catalog.

**Figure 9:**
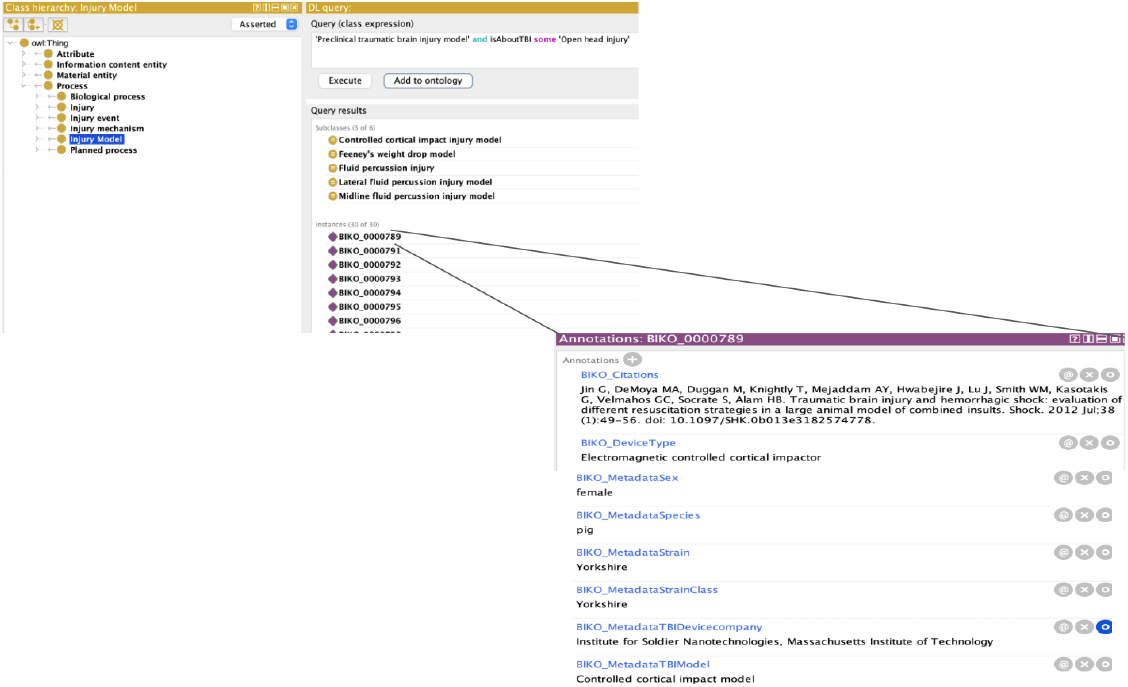
Use-case for BIKO with metadata from the PRECISE-TBI model catalog. Forty-two papers correctly classified as an open head injury based on the TBI model in each paper in the catalog from CQ1.

## Discussion

Here, we document the first version of BIKO, a machine-readable application ontology designed to standardize the reporting of methods, experimental parameters, and outcomes utilized in translational TBI research. The current version is tailored to address the specific needs of preclinical TBI research, with a future goal of adding clinical data to semantically bridge the divide between the bench and bedside. BIKO-TBI currently contains over 300 terms and 2000 statements about 4 major preclinical models relating explicit definitions of a preclinical TBI model with preclinical model attributes, phenotypes, and the measures used to assess them.

In order to create a formalized representation of preclinical TBI research, we made a few design choices. First, in BIKO, a TBI is caused by an injury event to the brain. To distinguish between preclinical TBI models and clinical TBI, the event is asserted to be either controlled, i.e., occurring in a lab through execution of a planned protocol, or uncontrolled - occurring in a natural environment, e.g., due to a fall at home, motor vehicle accident or sports injury. In either case, the result is a TBI in an organism. Each model results in some baseline phenotypes in the experimental subject that are relevant to the clinical condition, allowing researchers to retrieve models based on clinically meaningful phenotypes. For instance, as part of a controlled cortical impact model, a surgical procedure is used to remove a portion of the skull to expose the dura to mimic an open head injury.

We chose to model TBI as a process to reflect the evolution of injury over primary and secondary phases (Prins et al. 2013; Werner and Engelhard 2007). In fact, diseases and injuries are difficult concepts to model within ontologies from a realist perspective (Grenon and Smith 2004, Maynard et al. 2013). The BFO (Grenon, Smith, and Goldberg 2004), which is based on the realist perspective, divides entities into occurrents and continuants. A process is a subclass of occurrent. At its most basic level, an occurrent evolves through time while a continuant persists through time. In TBI, while the primary injury occurs at the moment of injury, it evolves over time to develop into a secondary injury or the secondary phase of the TBI. This second phase can last years after the initial injury, changing the profile of the injury. These temporal slices are often captured in disease classifications andcan rightly be considered as parts of temporal processes. On the other hand, one can look at a brain and point to the brain injury and measure its extent in many cases. A review of the 40 ontologies in in Bioportal of ontologies tagged as neurological disease (n=29) and neurological disorders (n=28) ontologies showed great variation in how disease or disorder was or was not classified. Of the 40 ontologies with the classification, they either: did not use BFO structure (n=27), did not have a disease or disorder as an explicit class (n= 3), were not in English (n=1), classified diseases as occurants (n= 2), classified diseases as disease continuants (n=4) and disease components were classified as occurants and continuants (n=3). Within the ontologies that used BFO structure, most ontologies only modeled diseases as continuants (67%) versus occurants (33%). Of interest, three ontologies had disease concepts components asserted as continuants and occurrents, namely the Alzheimer’s disease ontology (Malhotra et al. 2014), Neuropsychological Integrative Ontology (Gomez-Valades, Martinez-Tomas, and Rincon 2021) and the Epilepsy and Seizure Ontology (Sahoo et al. 2014). For instance, in the Alzheimer’s disease ontology the concept “Alzheimer Disease” was asserted as material entity (continuant), while the amyloid-beta deposition is a process. In a future version of BIKO, we are considering distinguishing between the injury course vs the injury itself, modeling the former as an occurrent and the latter as a continuant or disposition. In that way, the spatial extent of an injury such as might be measured histologically or in an MRI can be characterizes.

BIKOv1 focuses on modeling the basic preclinical concepts, ensuring flexibility in the design to allow for the extension of BIKO across preclinical and clinical studies. Future development will include deeper modeling of aspects of controlled injury that are known to differ across studies and to be clinically relevant, e.g., injury dynamic force. Additionally, future iterations will also include a formal way of describing more complex phenotypes associated with both the preclinical and clinical TBI and the intersection between the two. Finally, we will continue to utilize knowledge encoded in BIKO to provide semantics to individual or groups of preclinical CDEs (Kim et al. 2019; O’Connor et al. 2019). The semantics will serve as a bridge between the raw data and knowledge . For example, mapping data elements that comprise a behavioral test like the Barnes Maze to formal semantics can link the test to the types of functions it measures, e.g., memory. With significant efforts to improve rigor and reproducibility in TBI field many more potential use cases may materialize to further the development of BIKO, such as from initiatives like TRACK-TBI (Nelson et al. 2019; Meeuws et al. 2020), the Brain Trauma Blueprint TBI State of the Science (Pugh et al. 2021) development of precision diagnostics and therapeutics, ODC-TBI (Espin et al. 2019; Chou et al. 2022).

## Notes

### Competing Interest Statement

The authors have declared no competing interest.

